# De novo structural variants in autism spectrum disorder disrupt distal regulatory interactions of neuronal genes

**DOI:** 10.1101/2024.11.06.621353

**Authors:** Ketrin Gjoni, Xingjie Ren, Amanda Everitt, Yin Shen, Katherine S. Pollard

## Abstract

Three-dimensional genome organization plays a critical role in gene regulation, and disruptions can lead to developmental disorders by altering the contact between genes and their distal regulatory elements. Structural variants (SVs) can disturb local genome organization, such as the merging of topologically associating domains upon boundary deletion. Testing large numbers of SVs experimentally for their effects on chromatin structure and gene expression is time and cost prohibitive. To address this, we propose a computational approach to predict SV impacts on genome folding, which can help prioritize causal hypotheses for functional testing. We developed a weighted scoring method that measures chromatin contact changes specifically affecting regions of interest, such as regulatory elements or promoters, and implemented it in the SuPreMo-Akita software (Gjoni and Pollard 2024). With this tool, we ranked hundreds of de novo SVs (dnSVs) from autism spectrum disorder (ASD) individuals and their unaffected siblings based on predicted disruptions to nearby neuronal regulatory interactions. This revealed that putative cisregulatory element interactions (CREints) are more disrupted by dnSVs from ASD probands versus unaffected siblings. We prioritized candidate variants that disrupt ASD CREints and validated our top-ranked locus using isogenic excitatory neurons with and without the dnSV, confirming accurate predictions of disrupted chromatin contacts. This study establishes disrupted genome folding as a potential genetic mechanism in ASD and provides a general strategy for prioritizing variants predicted to disrupt regulatory interactions across tissues.

## Introduction

The human genome is folded into organized and hierarchical three-dimensional (3D) structures that span DNA loops, which bring two loci together, topologically associating domains (TADs), which insulate contact from surrounding regions, and compartments, which delineate regions of the genome based on activity. This complex multilayered system plays a critical role in regulating gene expression, in part by controlling the interactions between promoters and distal regulatory elements. Advancements in chromosome conformation capture technologies, such as HiC(1) and MicroC(2), have enabled the understanding of the interplay between the 3D genome and cellular function. Disruption of chromatin structures can lead to pathogenic re-wiring of regulatory interactions of disease-associated genes(3). For example, the fusion of two TADs can cause enhancer hijacking, whereby a gene in one TAD becomes regulated by a new enhancer and is differentially expressed. Furthermore, chromatin organization of neuronal cells plays a critical role in brain development and the onset of neurological disorders(4).

Structural variants (SVs)–a class of mutations encompassing large deletions, insertions, duplications, inversions, chromosomal rearrangements or combinations of these–have the potential to disrupt 3D genome structures and cause disease(5, 6). Germline SVs have been shown to cause gene misregulation and contribute to cancer and developmental diseases such as limb malformations(7), X-chromosome inactivation(8), limb morphogenesis(9), congenital malformations(10), branchiooculofacial syndrome(11), Fragile X syndrome(12), and Cooks syndrome(13). This evidence highlights the phenotypic consequences of disrupted genome structure in development and suggests the importance of further investigating SVs that alter genome folding as a potential causal mechanism in genetic disorders with poorly understood causes.

Since disrupted genome structure that leads to misexpression of critical genes has been characterized in many developmental disorders, we hypothesize that this mechanism is present in autism spectrum disorder (ASD). ASD is a class of neurodevelopmental conditions with complex causes that span environmental risk factors and genetics(14), with approximately equal contribution(15). While the heritability is estimated to be 40-80%(16), the genetic mechanisms are largely not understood with 80% of cases remaining without a genetic cause(17). Most studies that investigate the genetic causes of ASD focus on identifying common risk variants, mostly single nucleotide polymorphisms (SNPs)(18) or small insertions or deletions (indels). However, small sample sizes do not have enough statistical power to identify rare or low-risk variants, which can only be characterized if they fall on candidate risk genes(19, 20) but otherwise have largely unknown effects. Being able to evaluate the effect of rare variants is important because de novo mutations have been strongly implicated in ASD(20–23), and are estimated to contribute to about 10% of ASD cases(15). Furthermore, de novo variants are more likely to be causal in simplex families, in which only one offspring has been diagnosed with ASD. Some studies evaluated de novo variants using machine learning (ML) models to predict their effects on gene expression and found proband variants to be more damaging than those of siblings(24, 25). As this work did not include structural variants, there is a need for further evaluation of de novo structural variants (dnSVs) in ASD.

To evaluate ASD variants with unknown contribution to the disorder, we sought to test if they impact nearby neuronal developmental genes by disrupting their genomic contacts. Functionally characterizing variants requires individually evaluating their effect on gene expression and the accompanying mechanism, which is experimentally infeasible at scale due to time and cost limitations. To overcome this, ML models can be used with *in silico* mutagenesis (ISM) to predict variant deleteriousness at scale and prioritize candidate variants for experimental evaluations. One such model is Akita, a convolutional neural network (CNN) that predicts high resolution contact frequency maps from genomic sequence alone with very high accuracy(26). By making and comparing predictions for sequences with and without a mutation, ISM with Akita enables researchers to score its effect on 3D genome structure. For example, Akita accurately predicted how sequence variants that arose during hominid evolution altered genome folding in human versus chimpanzee neural progenitor cells(27). We previously developed SuPreMo-Akita (28), a computational pipeline that streamlines ISM with Akita allowing us to test massive numbers of rare and complex variants for their effects on genome structure, prioritizing variants and generating testable hypotheses about their effects on disease-relevant genes.

However, SuPreMo-Akita does not specifically identify changes in genome folding that affect gene regulatory interactions. To address this limitation, we developed a weighted scoring method and added it to the SuPreMo software package. Then, we leveraged this functionality in combination with excitatory neuron (ExN) PLAC-Seq data(29) to evaluate the effects of dnSVs at neuronal putative cis-regulatory element interactions (CREints). We found that dnSVs present in ASD individuals(30) are more disruptive to CREints than dnSVs in unaffected sibling controls. We defined a set of criteria to prioritize variants that are likely to be causal and can feasibly be tested in ExN cells. Using CRISPR-engineered induced pluripotent stem cell (iPSC)-derived ExNs, we tested a 23 kilobase (kb) deletion in a region rich in neuronal genes. The results show a striking similarity between experimental and computationally sredicted changes in genome folding and detected differential expression of many neurodevelopmental genes. This provides a proof-of-concept for testing variants of unknown significance for their effects on chromatin interactions.

## Results

### Large ASD dnSVs disrupt genome structure via CTCF binding sites

In this study, we used dnSVs from simplex families in the Simons Foundation Autism Research Initiative (SFARI) Simons Simplex Collection (SSC) cohort(30, 31). Variants from ASD probands (n=521) and their unaffected siblings (n=348) were previously called from short-read whole genome sequencing(30). We scored these variants for their predicted effects on 3D genomic contacts in the surrounding region using previously published SuPreMo-Akita(28) (**Fig. 1A, Methods**). As a part of this pipeline, contact frequency maps that correspond to the reference and alternate allele and the surrounding region are compared using Spearman’s correlation (hereafter referred to as correlation). The resulting score (1 - correlation) is a measure of how disruptive each variant is to genomic contacts in the surrounding ∼1 Mb region. Disruption scores were generated for a subset of dnSVs that are compatible with SuPreMo-Akita based on their length, type, and region (**Methods**). Across the 587 scored variants, the mean disruption score was higher for proband versus sibling dnSVs (**Fig. 1B, S1B**). The disruption score distributions for both groups are left skewed with most variants resulting in low disruption to genome folding, consistent with other studies generating ISM disruption scores with Akita(32, 33). But the highest scoring variants include more proband than sibling dnSVs (14% and 7% of each group have 1 - correlation > 0.2, respectively), suggesting that the difference in means is driven by high scoring variants.

**Figure 1.**
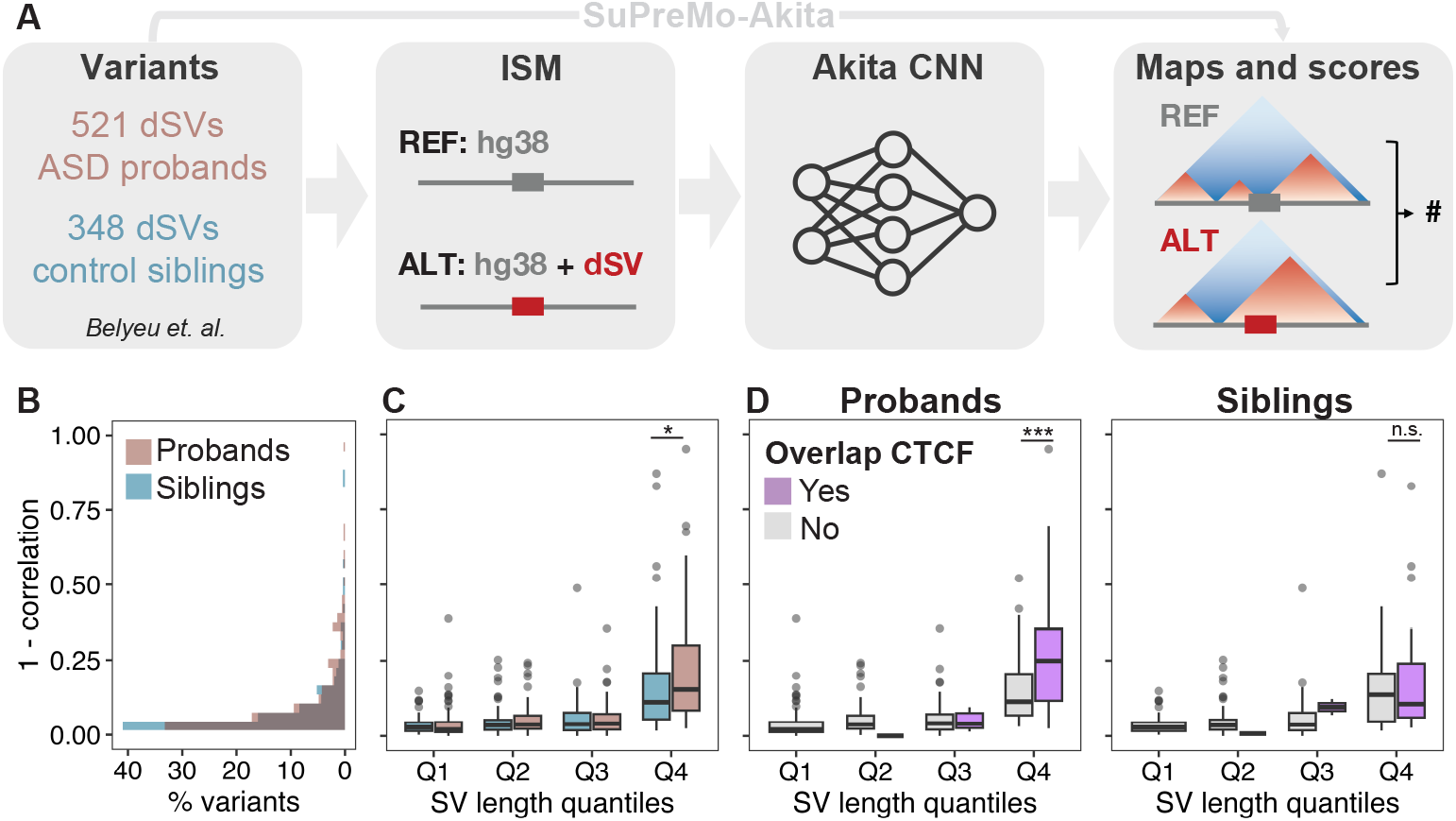
ASD dnSVs are predicted to be more disruptive to 3D genome folding than controls. **A**. SuPreMo-Akita workflow for scoring variant disruption to genome folding. Variant information is inputted into SuPreMo-Akita, which in turn generates reference and alternate sequence pairs and inputs those into Akita. It then processes the resulting maps and compares them to generate disruption scores. **B**. Distribution of disruption scores for dnSVs from probands (pink) and siblings (blue). Mann-Whitney U test p-value is 0.039. **C**. Disruption scores across variant length quantiles, Q1-Q4, for proband and sibling dnSVs. Length quantile cutoffs are 147 bp, 3,976 bp, and 32,549 bp. Number of dnSVs per quantile are as follows: 78 and 72 in Q1, 77 and 73 in Q2, 97 and 51 in Q3, and 97 and 53 in Q4 for probands and siblings, respectively. Mann-Whitney U test p-value for Q4 is 0.032. **D**. Disruption scores across length quantiles for probands (left) and sibling (right) dnSVs separated by whether the variant coordinates overlap at least one CTCF binding site (purple) or not (gray) using ChIP-Seq data from ExNs. Mann-Whitney U test p-value for Q4 is 0.002 and 0.630 for proband and sibling dnSVs, respectively.

In order to understand factors that contribute to high scoring proband dnSVs, we looked at how scores relate to variant length, its type, and functional features that it overlaps. We find that dnSV length is positively correlated with disruption scores, with longer variants causing bigger changes to genomic contacts (**SI Appendix, Fig. S1C**). We therefore grouped variants by their length quantiles, to ensure the differences between proband and sibling dnSVs are not solely due to length differences. We find that proband dnSVs are significantly more disruptive than sibling dnSVs when controlling for dnSV length, but only in the fourth length quantile (**Fig. 1C**). This further supports the idea that large variants are driving these differences, while smaller variants are equally damaging. Next, we evaluated how disruption scores compare across variant types and found significant differences, namely that duplications are the most disruptive and inversions the least (**SI Appendix, Fig. S1D**). This trend could be partially explained by dnSV length since duplications are also the largest variants in this dataset, while inversions are the smallest (**SI Appendix, Fig. S1E**). Nonetheless, these trends do not contribute to the difference in scores between proband and sibling dnSVs, because the two groups have similar distributions of variant types (**SI Appendix, Fig. S1F-G**).

Lastly, we evaluated how variant overlap with CTCF binding sites relates to disruption score. Given that CTCF plays a key role in determining genome structure through cohesin-mediated loop extrusion, we expected variants that disrupt CTCF binding to result in larger changes to genome folding. We see this pattern across both proband and sibling dnSVs, although the association is stronger in probands (**SI Appendix, Fig. S1G**). When categorizing dnSVs by length quantiles, we find that only the longest dnSVs are significantly more disruptive when overlapping CTCF, and this is only true in probands (**Fig. 1D**). This suggests that proband dnSVs disrupt CTCF binding sites that are more important for genome folding and result in more consequential changes when disrupted compared to sibling dnSVs. Overall, these results show that proband dnSVs are slightly but significantly more damaging to chromatin organization than the unaffected sibling dnSVs, suggesting that disrupted genome folding might play a role in the genetic etiology of ASD.

### ASD dnSVs disrupt neighboring neuronal regulatory element contacts

To focus our scoring approach on variants where the altered chromatin contacts affect regulatory elements and their target genes, we defined putative cis-regulatory interactions (CREints) as chromatin interactions that are within the ∼1 Mb prediction window, and correspond to the promoter of an expressed gene (**SI Appendix, Fig. S2A, Methods**). To apply this approach to ASD, we used H3K4me3 PLAC-Seq and RNA-Seq data from primary ExNs(29)–the cell type most implicated in the disorder(34). We observed that proband but not sibling noncoding dnSVs are enriched near CREints (**Fig. S2B**). This suggests that while these variants do not directly impact any coding sequences, they have potential to alter regulatory interactions of genes expressed in ExNs.

Motivated by this observation, we sought to specifically evaluate the effect of dnSVs on gene regulation by generating scores that only reflect disrupted interactions at CREint anchors (putative promoter and regulatory element). We developed a weighted scoring method which is flexible to various annotation types and can be tuned to emphasize regions of interest (ROIs) while still considering changes within the locus (**Fig. 2A, Methods**). Instead of comparing the maps as a whole, we calculated disruption scores by averaging the disruption at each bin in the prediction window (**Fig. 2A:** disruption track in purple). This allows for calculating the weighted average by multiplying the disruption track by an ROI weight track before taking the mean (**Fig. 2A:** weight track in orange). The weight track upscales disruptions at bins corresponding to ROIs by a user-specified amount. Weighted disruption scores are lower than unweighted scores when there is less disruption at ROI bins versus the rest of the map (**SI Appendix, Fig. S2Ci**), greater than unweighted scores when there is more disruption at ROI bins (**SI Appendix, Fig. S2Cii**), and similar to unweighted scores when ROI and non-ROI bins have similar disruption values (**SI Appendix, Fig. S2Ciii**).

**Figure 2.**
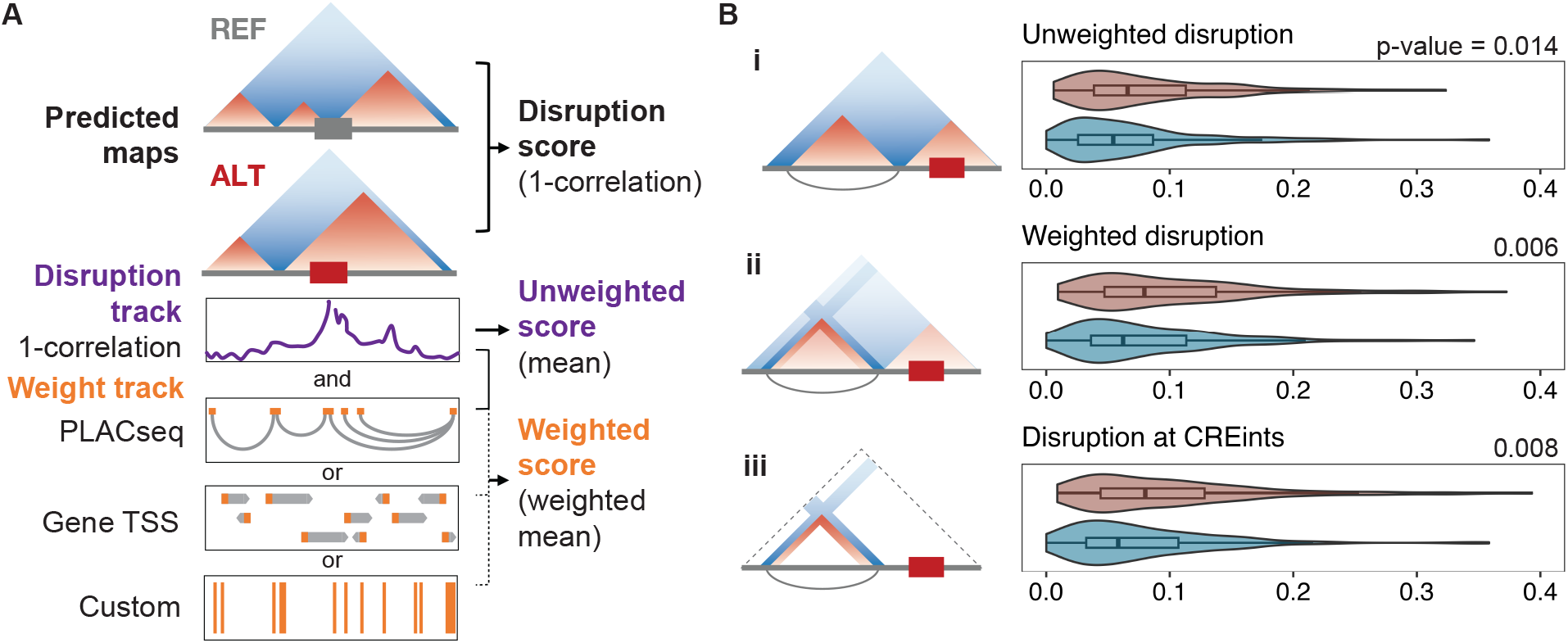
Neuronal regulatory element interactions are indirectly disrupted by variants in ASD. **A**. Schematic of SuPreMo-Akita implementation of weighted scoring. Unweighted score is the mean of the disruption track (purple). For the weighted score, each value in the disruption track is multiplied by the corresponding value of a weight track (orange, several options shown) before taking the average for the window. The weight track may be generated from PLAC-Seq paired regions, custom input regions, or transcription start sites (TSSs). If PLAC-Seq data is inputted, it will be processed to filter and condense loops (**Methods, SI Appendix, Fig. S2A**). **B**. Disruption scores for a subset of dnSVs that are close enough to CREints to both be in the same Akita prediction window. Scores, from top to bottom, correspond to: (i) original scores where the dnSV is centered in the sequences, (ii) scores where the dnSV and its neighboring CREInt are centered in the sequences, (iii) scores for windows in (ii) where CREint anchors are upweighted 10-fold, (iv) scores for windows in (ii) using only bins at CREint anchors.

To easily and scalably apply weighted scoring to the ASD dnSV dataset, and to make it accessible for others working with models beyond Akita, we incorporated the method into the SuPreMo pipeline. In sum, we added inputs for: ROIs and the scaling factor, the Akita-prediction cell type, and the extent to shift the prediction window for each variant to include ROIs. To input ROIs and scaling factors, arguments ‘--roi’ and ‘--scale’ can be used to provide regions to upweight and the amount to upweight by, respectively. These ROIs can be PLAC-Seq paired regions, custom regions, or TSSs for example (**Fig 2A, Methods**). To include neighboring ROIs in the prediction window, the ‘--shifts’ parameter can be used to specify the up- or down-stream shifting amount for each variant. This shifting feature also makes SuPreMo compatible with models, like ExPecto(35), that require centering a TSS instead of the variant. The adapted SuPreMo pipeline with all these modifications was released as version 2 (V2) on GitHub (**Methods**).

With this tool in hand, we turned back to evaluating dnSVs likely to change chromatin contacts that affect neuronal CREints in ASD probands and unaffected siblings. We focused on the 319 dnSVs that are near at least one CREint. Our original scores did not show significant differences between probands and siblings for this subset of dnSVs (Mann-Whitney U test p-value 0.36). We then paired each dnSV with all the nearby CREints and used custom shifting to make predictions centering each dnSV and CREint pair (**Methods**). We calculated unweighted and weighted disruption scores for each pair and kept the highest score for each dnSV. We used all of the CREints in the window for weighting, but found that using only the paired CREint for each window did not change the trends we observed. While we see a significant difference between proband and sibling dnSV unweighted scores, weighted scores enhance this difference both when up-weighting CREints by 10-fold (**Fig. 2Bii**) and when only scoring bins at CREint anchors (**Fig. 2Biii**). Additionally, there are more proband dnSVs with higher weighted versus unweighted scores (50% of dnSVs) compared to siblings (36%) (**Fig S2D-E**). This data suggests that proband dnSVs are more disruptive at CREints and are therefore more likely to have a functional impact in ExNs. For the majority of variants, if scores at CREints were higher than unweighted scores, CREint scores were the highest, further supporting that changes are concentrated at CREint anchors (**Fig S2E**). In summary, we found that proband dnSVs are more disruptive to neuronal promoters and their distal regulatory elements as compared to sibling dnSVs and that predicted changes in chromatin contacts in disrupted loci are concentrated on the regulatory interactions. These results further support our hypothesis that proband dnSVs contribute to ASD partially through 3D genome folding disruption.

### Prioritizing candidate dnSVs likely to affect ASD genes

Our SuPreMo-Akita weighted disruption scores produced a testable hypothesis for each dnSV about what regulatory interactions it may alter in neurons. Genome editing and induced pluripotent stem cells provide experimental tools for investigating these hypotheses. Since generating stable, isogenic cell lines is currently relatively low throughput, we sought to prioritize a small number of dnSVs with compelling evidence in support of functional characterization. To do so, we compiled information about the variant, the prediction, and the proband, organizing these data into 7 required and 5 optional criteria. These criteria prioritize disruptive dnSVs near–but not overlapping–ASD genes, have robust Akita predictions, and occur in probands without a high risk variant elsewhere in their genome (**Methods, SI Appendix, Fig. S3A**). Most dnSVs meet 4-5 of the required criteria, and typically proband dnSVs meet more criteria than sibling dnSVs (**SI Appendix, Fig. S3B**). Nine dnSVs passed all 7 required criteria and were evaluated both on the 5 optional criteria and on qualitative characteristics, such as visually inspecting how strong the changes are, where they fall with respect to ASD genes, and how PLAC-Seq loops support the prediction (**Methods**).

With this selection process, we identified three proband dnSVs that have the potential to contribute to ASD by misregulating associated genes through 3D genome folding disruption (**SI Appendix, Fig. S3C**). First, a 160 bp deletion in chromosome 17 (**SI Appendix, Fig. S3Ci**) lowers contact frequency at and near RAI1 (arrows), a high confidence and syndromic SFARI gene(36). RAI1 is a transcriptional regulator of the circadian clock involved in embryonic development and neural differentiation(37). In another proband, a 2 Mb duplication encompasses RAI1 further supporting its association with ASD. This deletion overlaps a MYO15A exon, but the gene is lowly expressed in ExN and not relevant in ASD. Second, a 1.6 kb deletion in chromosome 9 (**SI Appendix, Fig. S3Cii**) results in slightly decreased contacts of STXBP1 (arrows), a high confidence SFARI candidate involved in 49 ASD cases that plays a role in neurotransmitter release(36). Of note, this variant was found in a proband that also has a missense mutation in an exon of SNX14, a syndromic SFARI gene, which might contribute to their genetic cause of ASD. This is an intronic deletion in NIBAN2 and overlaps two ExN H3K27Ac peaks. Lastly, a 23 kb deletion in chromosome 10 (**SI Appendix, Fig. S3Ciii**) lies in a locus rich with neurodevelopmental genes, including PPP3CB, which is downregulated in two in vivo ASD models(38, 39), CAMK2G, which is linked to neurodevelopmental disorders and associated with ASD, and P4HA1, which was found to be associated with ASD in a de novo risk score analysis(22). The deletion overlaps 8 exons of USP54, which is not involved in ASD or neurodevelopment and has a very low probability of loss of function (pLI = 0) suggesting this heterozygous deletion may not affect USP54 protein function. While the deletion does not overlap any ExN regulatory marks, it removes a CTCF binding site (**SI Appendix, Fig. S4A**) suggesting that the mechanism behind the changed contact maps includes the loss of an insulating boundary. Indeed, disruption scores from tiled 1bp deletions across the dnSV recapitulate the CTCF motif (**SI Appendix, Fig. S4B**), also highlighting Akita’s ability to capture the grammar of genomic determinants of 3D genome folding. Together, these features made this chromosome 10 variant the strongest candidate of the five and prompted us to experimentally test the predictions.

### In vitro variant model validates predictions and suggests transcriptomic impact

Since regulatory interactions are likely to vary across cell types, neuronal chromatin folding patterns are different from non-neuronal cells(4), and ASD largely affects ExNs, we characterized the chromosome 10 proband dnSV in WTC11 i^3^N iPSC-induced ExNs (38). Using a CRISPR/Cas9-mediated deletion in the WTC11 i^3^N iPSC line(40), we generated two clonal cell lines with a homozygous deletion similar to the variant (“full deletion”; **SI Appendix, Fig. S4A, Methods, SI Appendix, Table S1**), allowing us to isolate the effects of the deletion on genome folding without being influenced by an unedited copy. In contrast to the variant in the proband, which affects exons 11-18, is in frame, and should result in a partial mRNA expression of 14 exons of USP54, the engineered clones harbor a deletion of part of exon 13 and exons 14-18, and result in an early stop codon with in frame transcription resuming after the deletion, including exons 19-22 (**SI Appendix, Fig. S4B-C**). USP54 is involved in ubiquitin-proteasome-dependent proteolysis and is expressed in primary ExNs (TPM = 16)(29). We differentiated two unedited (WT) and the two isogenic full deletion clones into ExNs and performed HiC to evaluate chromatin interactions (**Methods**). The contact frequency map from WT ExNs closely matched the Akita prediction from the reference human genome, and the observed effect of the deletion on the map was highly similar to our prediction for the proband variant (**Fig. 3**). Namely, the overall strengthened contact (purple) between the TAD boundaries, the decreased contact (green) of the upstream boundary with inter-TAD regions (left arrow), and increased contact of the downstream boundary with inter-TAD regions (right arrow), are consistent between the prediction and experimental HiC data. To test whether the predicted changes were due to the deleted CTCF binding site specifically, we generated two additional clonal cell lines in which only the CTCF ChIP-Seq peak was deleted (“CTCF deletion”; **SI Appendix, Fig. S4A-C**). Both reference and CTCF deletion contact frequency maps closely matched Akita predictions, resulting in a similar, but less strong, effect as the full deletion (**SI Appendix, Fig. S5B-C**). The more striking changes caused by the full deletion suggest that while the CTCF binding site plays a role in chromatin interactions in this locus, additional sequences within the full ASD dnSV likely also contribute.

**Figure 3.**
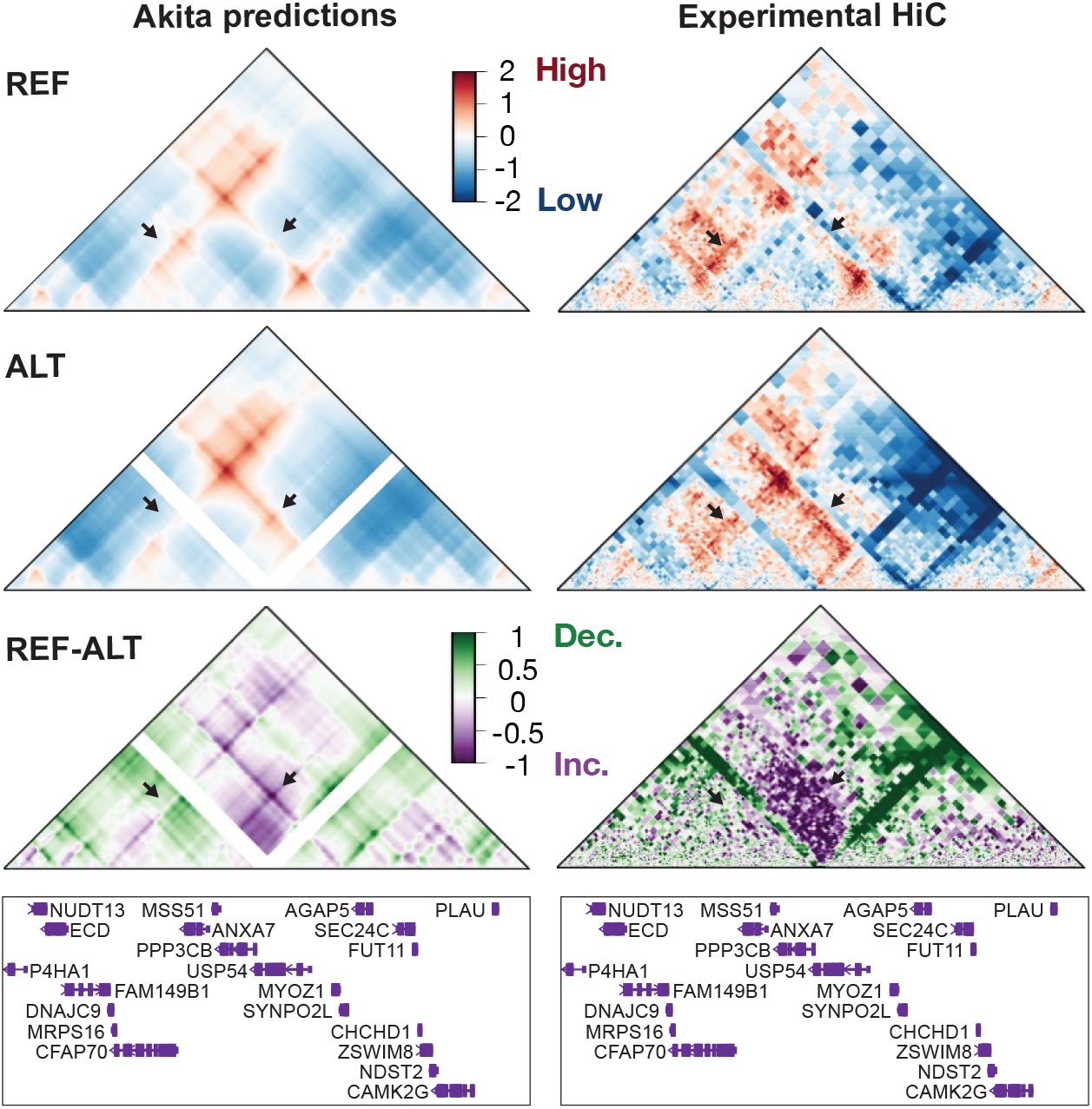
Convolutional neural network correctly predicted effect of an ASD deletion on chromatin contacts. Predicted (left) and experimental (right) contact frequency maps for the reference genome (top), reference genome with the 23-kb chromosome 10 deletion (middle) and the difference between the two (bottom). Genes in the corresponding ∼917 kb region shown below maps. Arrows point to two regions with changes in contact.

To evaluate the effect of the chromosome 10 deletion on gene expression, we performed RNA-Seq on cell lines from all three conditions (2 WT, 2 full deletion, 2 CTCF deletion). We first evaluated the effect of the deletions on USP54 and found that its transcription continued after both deletion types (**SI Appendix, Fig. S6A**). A principal component analysis (PCA) of the gene expression data clearly separated all three editing conditions (**SI Appendix, Fig. S6B**). It is unclear how much of the separation between the full deletion and CTCF deletion cell lines is due to the strength of the variant-caused contact change or the differing effect on USP54. We focus our next analyses on the full deletion since it is similar to the variant found in the proband and the in vitro changes are most likely to mirror its expected effect. We then performed a genome-wide differential gene expression analysis and found that the full deletion resulted in 1,102 differentially expressed genes (DEGs) (**SI Appendix, Fig. S6C, SI Appendix and Dataset S1**). Interestingly, full deletion DEGs are enriched for GO terms involving neuron development, differentiation, and function, when using all expressed genes as a background (**SI Appendix, Fig. S6D and Dataset S2**). They are also enriched for ASD genes (**SI Appendix, Fig. S6E-F**). Though the damaged USP54 transcript likely imparts some differential expression signal, we do not see enrichment of genes from the Ubiquitin-Proteasome Dependent Proteolysis pathway in either deletion cell line (**Fig S6E-F**), suggesting it is not a dominant driver. Overall, the transcriptomic effects of the chromosome 10 deletion cell model further support its potential to affect the function of neuronal cells.

We speculate that changed genomic contacts also play a role in differential expression by misregulating genes near the deletion and resulting in a cascading effect genome-wide. Interestingly, the full deletion did not affect the expression of genes in the ∼1 Mb Akita prediction window; however, HiC data allows us to evaluate the effects of the variant on genes past this narrow window. These long-distance effects include regions with important neurodevelopment genes such as DDIT4, which is 1.2 Mb from the dnSV and has potential to be affected by changed genomic contacts (**SI Appendix, Fig. S7A**). DDIT4 is involved in neuron differentiation and death and has been implicated in autism in two different studies which found it was downregulated in ASD(41, 42). The gene is also significantly downregulated in the full deletion cell line (**SI Appendix, Fig. S7B**). DDIT4 is a part of the mTOR signaling pathway, genes from which are enriched in the full deletion DEGs (**SI Appendix, Fig. S7C**, odds ratio = 6; chi-squared p-value = 2.2e-14), suggesting that a subset of DEGs is caused by DDIT4 downregulation. The CTCF deletion cell lines also showed similar contact changes at DDIT4 and decreased expression although both these changes are less pronounced than in the full deletion (**SI Appendix, Fig. S7A-B**). While the role of the 3D genome folding changes caused by the full deletion on gene expression is unclear, this data supports a scenario where the variant impacts DDIT4 expression by disrupting its distal regulatory contacts and in turn misregulates an array of genes in its pathway. Other nearby DEGs that could have a similar effect are SPOCK2, which is thought to be involved in neurogenesis, and UNC5B, which is involved in embryogenesis and nervous system development. More work needs to be done to link the differences in genomic contact to differences in gene expression and ultimately to ASD.

## Discussion

Here, we assess the impact of ASD-associated structural variants on genome organization and introduce a pipeline for prioritizing variants that may contribute to the disorder by disrupting distal regulatory interactions. We enhance the SuPreMo-Akita pipeline with two new features to evaluate the effects of variants on specific nearby regions of interest. This software tool and our proof-of-principle analyses will enable variant prioritization for other cohorts and biological contexts. Our analyses of ASD probands and their unaffected siblings revealed chromatin interactions are more often altered by ASD dnSVs, particularly those involving promoters of neuronal genes. Notably, we identified several candidate ASD de novo deletions predicted to affect genome folding near risk genes. We validated one of our predictions using CRISPR genome editing and iPSC-derived ExN cells, demonstrating that a 23-kb deletion overlapping a CTCF binding site increases contacts between previously insulated genomic regions. This deletion results in misregulation of neurodevelopmental genes across the genome and may be implicated in the proband’s diagnosis, although further studies are needed to confirm this.

Finding appropriate controls for structural variants–such as random length-matched SVs or ones from healthy individuals–is challenging due to their likelihood of falling in protected regions or being generally smaller. Studying variants in simplex families is especially valuable because the unaffected siblings share the same genetic background as the proband, enhancing the relevance of our comparisons. However, siblings are still more likely to possess damaging variants compared to unrelated individuals, reflecting the genetic complexity of ASD and the challenges in diagnosis. Indeed, we found examples of sibling dnSVs that are predicted to disrupt contacts of neuronal genes, even though they may not be causal for ASD. Consequently, we did not expect drastic differences in variant effects between probands and siblings. Despite these challenges, the ability to detect significant differences in disruption scores is encouraging and underscores the robustness of our findings. This is the first evidence at a cohort scale that disrupted genome folding may play a role in ASD.

Studying large structural variants poses significant challenges, particularly when it comes to experimental modeling. Recapitulating large heterozygous variants, such as the 23 kb deletion on chromosome 10, is constrained by factors like PAM site availability, gRNA efficiency, and allelic differences. The WTC11 cell line lacks sufficient heterozygous SNPs for allele-specific HiC analysis that would mirror Akita’s allele agnostic predictions. To address these issues, we introduced a homozygous deletion. Although this approach complicates the interpretation of gene expression changes, it effectively isolates and confirms the impact of the deletion on genomic contacts. Due to gRNA efficiency limitations, the CRISPR-induced deletion results in an early stop codon that disrupts transcript continuity, differing from the in-frame deletion present in the proband. Despite these limitations, the generated clonal cell line closely models the variant’s effects in neurons. Because USP54 is not a known ASD or neurodevelopmental gene, and it is not functionally associated with the DEGs we detected, we hypothesize that USP54 is a bystander gene and that the DEGs result from downregulation of DDIT4 and/or other neuronal genes affected by the dnSV. But further experimental studies are needed to confirm this. Future work focused on engineering a more precise heterozygous deletion could provide further insights into gene expression changes and help identify potential functional phenotypes.

This work introduces a pipeline for assessing variant causality and prioritizing candidate disease variants. Its application to ASD demonstrates that variant-induced alterations in genome folding may contribute to the disorder and identifies a candidate causal deletion. Teasing apart ASD’s complex etiology will require a holistic understanding of variant effects. This may include new genetic underpinnings, which our study aims to predict with ML. Our study aims to clarify the genetic underpinnings of such disorders by individually characterizing variant effects with the SuPreMo-Akita workflow. Our updated software enables this strategy to be efficiently applied to other genetic diseases and biological contexts to generate and prioritize testable hypotheses about potentially causal variants. Future work could enhance SuPreMo’s utility by integrating it with other predictive models and annotating variants based on their impact on epigenetic modifications and gene expression. This ISM pipeline has the potential to uncover new disease-associated variants for experimental validation, increasing the yield of genome editing studies and advancing our understanding of the molecular mechanisms underlying a wide array of genetic disorders.

## Materials and Methods

### Previously published data sources

SFARI SSC proband and sibling dnSVs were previously called from short-read whole genome sequencing data using the GATK-SV pipeline (30, 43). PLAC-Seq data is from primary human mid-gestational ExNs(29). ExN epigenetic data was downloaded from publicly available sources. ChIP-Seq data is from ENCODE with the following dataset IDs: ENCSR489QDF for CTCF, ENCSR201NXK for H3K27me3, ENCSR818PER for H3K27Ac, and ENCSR411ZUA for H3K27me3. Neural progenitor cell (NPC) HiC data was previously published(27). ASD genes are cataloged from three different datasets: SFARI genes(36), risk genes from an exome sequencing study(23), and genes found from de novo risk scores(22). Gene exon coordinates used are from GENCODE v46(44).

### Variant scores

To measure the predicted disruption on nearby chromatin contacts across 869 SVs, SuPreMo-Akita was used. While Akita was trained on 5 different cell types, none of these was a neuronal cell type. We used predictions for Human Foreskin Fibroblast (HFF) cells since this cell type had the best performance and Akita’s predictions for all five cell types were highly similar(26). For each SV, this pipeline takes the 1 MB reference sequence surrounding the variant (center coordinate) and generates a 1 MB mutated sequence with the alternate allele of the variant changed at the appropriate location. Both sequences are input to the Akita model and the resulting contact frequency maps are padded and masked so that each 2,048 bp bin from either map corresponds to a similar sequence. The remaining regions of the maps–those outside of the SV itself–are then compared using correlation and mean squared error (MSE). To get augmented scores, scores from four sequences were averaged: 1) no augmentation, 2) 1 base pair (bp) right shift, 3) 1 bp left shift, and 4) the reverse complement. Of note, differences in scores across variant types could be partially due to how SuPreMo-Akita scores different variant types, for instance duplications are treated like insertions of the duplicated sequence at its 5’ end(28).

### Variant exclusions

Only a subset of dnSVs were scored–they have to be above the length threshold, have known alternate allele sequences (deletions, inversions, duplications, and complex variants), and be in regions where the reference sequence is known. Scores represent changes in the region surrounding each variant, excluding the variant itself. As a result, the size of these regions varies depending on the size of the variant. This could overestimate the impact of very large variants, so we did not evaluate 73 variants larger than 700 kb. These variants could be tested with genome folding prediction models that use larger input sequences, such as ORCA(31), however the resulting predictions would be at much lower resolution and would preclude analysis of individual CREInts. Variants for which the alternate allele sequence is not known were excluded from the analyses, because Akita needs the exact sequence to generate a prediction. This includes insertions (SV types: ALU, LINE1, SVA, and INS) and translocations (SV type: CTX). SuPreMo filters out variants that are greater in length than ⅔ of the input sequence. Additionally, certain variants do not get a score because the region that they are in has a composition of unknown sequence of greater than 5%. Since CPX variants are not compatible with SuPreMo, we wrote custom code to generate mutated sequences with those variants. However, these variant types are excluded from the CREint-weighted analyses (**Figures 2 and S2**), since those analyses require using the SuPreMo pipeline. In total, we generated scores for 598 dnSVs.

### Neuronal cis regulatory element interactions

To annotate CREints, paired peak regions from primary human ExN PLAC-Seq data were used(29). Here, we refer to each pair of regions as a loop and each single region as a loop anchor. Loops were processed by grouping and filtering to generate regions of interest (ROIs) for weighted scoring (**SI Appendix, Fig. S2A**). Only interchromosomal loops with anchors closer than 900 kb were considered, as limited by Akita, which outputs contact frequency maps corresponding to a 917,504 bp region. Then loops within 10 kb of each other were considered redundant and grouped together to result in less pairs and better fit the Akita 2,048 bp bin resolution. For all the pairs with the last left anchor, the right anchors that were within 10 kb were combined so that the new right anchor for all the pairs is the same and includes all the previous right anchors. The same process was done on the left anchors of pairs with the same right anchor. While all of these steps are also a part of how SuPreMo processes inputted regions of interest for weighting, in this study we also took additional filtering steps. We only kept CREints at least one anchor of which overlaps the promoter–defined as the 2kb outside of the TSS–of genes expressed in ExNs(29). Lastly, after pairing CREints with variants, we only kept ones where both loops of the CREint and the variant are less than 900 kb and therefore fit in the prediction window. Note that any plotted CREints are not subject to this filtering but rather the initial PLAC-Seq pairs. While CREints may include non-regulatory contacts and introduce false positives (FP), the FP rate will be consistent between probands and siblings, allowing for their fair comparison.

To calculate enrichment of dnSVs near CREints, the genome was split into 1 Mb bins and each bin was annotated for whether it overlapped with a dnSV and a CREint region. A chi-squared test was used to calculate the p-value.

### CREint weighted scoring

For weighting scores at ROIs, contact frequency maps were generated as described above. Instead of comparing the whole map, disruption tracks were generated by comparing each 1-bin-wide column in the map and getting a list of 448 scores across all bins. Then a weight track was generated, which has a scalar value in bins that overlap ROI and 1 in the rest of the bins. The mean of the disruption track is the unweighted score and the mean of the disruption track multiplied times the weight track is the weighted score. The weight track is generated by upscaling any bin that overlaps any ROI. ROIs can be given in the form of a bed file for a custom set of regions, such as accessible regions from ATAC-seq, a paired bed file, such as promoter contacts from PLAC-Seq, or by just specifying ‘genes’ which will use a 2 kb region centered at the TSS of every gene.

To find variants near CREints, CREints and variants were paired together if both CREint anchors and the variant were all with 900 kb. Pairs where the variant overlapped either CREint anchor were removed. The resulting 3,677 pairs were made up of 347 dnSVs and 2916 CREints. Disruption scores for this subset of dnSVs were extracted from the existing set of scores. Then, for each pair, a shift for the prediction window was calculated that would result in the pair being centered in the Akita input sequence. These shifted windows were scored using SuPreMo and 10-fold up-weighting either all CREint anchors in the window or only anchors of the paired CREint. This resulted in a score for each dnSV-CREint pair which was summarized into one score per dnSV by taking the maximum across all CREint pairs.

### Criteria for variant prioritization

We used the following criteria for prioritizing candidate dnSVs.

Required

1. **Not on ASD gene:** Variant does not overlap ASD gene exon.
2. **No causal dnSV:** Proband does not have dnSV overlapping ASD gene exon.
3. **Good prediction:** Predicted maps for the reference genome around dnSV are similar to NPC experimental HiC maps. This includes dnSVs for which the mean squared error (MSE) between the Akita predicted and the experimental contact maps is less than the 85th percentile of MSE across all scored variants.
4. **No sibling dnSV:** There are no similar variants in siblings. This includes proband dnSVs that don’t overlap by more than half of the variant region with more than half of any sibling dnSV region.
5. **Disrupts CREint:** Variant results in changed contact at CREints. This includes dnSVs with weighted disruption scores at CREints (**Fig. 2Biv**) larger than the 65th percentile across all standard scores (**Fig. 1B**).
6. **Near ASD gene:** Variant is within 500 kb of an ASD gene.
7. **Disruptive:** Variant is disruptive to 3D genome folding of surrounding regions. This includes dnSVs with disruption scores above the 65th percentile across all scores. Optional:
8. **Not on expressed gene:** Variant does not overlap expressed gene (TPM > 0.5) exon.
9. **Not on ExN RE:** Variant does not overlap ExN regulatory elements, namely active enhancers (H3K27Ac), poised enhancers (H3K4me1), and poised promoters (H3K27me3) as defined by their respective ChIP-Seq peaks.
10. **Change on CREint:** Variant disruption focused on CREints, meaning weighted score (**Fig. 2Biii**) is higher than unweighted score (**Fig. 2Bii**).
11. **On CTCF:** Variant overlaps less than half of any ExN CTCF ChIP-Seq peak.
12. **Deletion:** Variant is a deletion. These are the most straightforward to edit in cells.

Variants that passed all 7 required criteria were qualitatively evaluated. The selected three deletions (**SI Appendix, Fig. S3**) were predicted to cause structural changes to the chromatin structure, as opposed to just changes in contrast. The changed contacts align with nearby ASD genes and excitatory neuron PLAC-Seq loops correspond well with Akita-predicted contacts. Additionally, the deletion that was selected for experiments had the most reproducible effect when augmenting maps with sequence shifting and reverse complement.

### CRISPR-engineered cell lines

The isogenic clonal iPSC lines were made by CRISPR/Cas9 system-mediated deletion. The WTC11 i^3^N iPSC line with doxycycline-inducible Ngn2 integrated at the AAVS1 safe harbor locus was used as the parental line. We designed four sgRNAs using CHOPCHOP (https://chopchop.cbu.uib.no/) to delete the genomic region overlapping with the ASD variant and a CTCF site within the ASD variant. The designed sgRNAs (**SI Appendix, Table S1**) were in vitro transcribed using the Precision gRNA Synthesis Kit (Invitrogen, A29377), and Cas9-NLS protein was ordered from QB3 MacroLab at the University of California, Berkeley. We assembled the Cas9/sgRNA complex by incubating the in vitro transcribed sgRNAs and Cas9-NLS protein at 20-25°C for 15 min and delivered the complex into WTC11 i3N iPSCs using nucleofection (Lonza, VPH-5012). After nucleofection, the cells were seeded into Matrigel-coated (Corning, 354277) wells for recovery. 3-4 days later, we sorted live cells into 96-well plates with one cell per well using fluorescence-activated cell sorting (FACS) to generate clonal cell lines. About two weeks later, the viable clones were expanded. Meanwhile, we extracted genomic DNA from each clone using QuickExtract DNA Extraction Solution (Biosearch Technologies, QE09050) and checked the genotype of each clone using PCR and Sanger sequencing. This resulted in 6 clonal cell lines included in this manuscript: 2 WT, 2 full deletion, and 2 CTCF deletion, each a pair of biological replicates.

### ExN differentiation

The WTC11 i3N iPSCs were cultured on Matrigel-coated (Corning, 354277) plates and maintained in mTeSR Plus media (STEMCELL Technologies, 100-0276), and passaged with Accutase (STEMCELL Technologies, 07920) and 10-μM ROCK inhibitor Y-27632 (STEMCELL Technologies, 72302). The cells were grown with 5% CO_2_ at 37°C and verified mycoplasma-free using the MycoAlert Mycoplasma Detection Kit (Lonza, LT07-218). We differentiated the iPSCs into ExNs by using a two-step differentiation protocol. First, we cultured iPSCs on Matrigel-coated plates with pre-differentiation media with doxycycline (2 µg/mL; Sigma-Aldrich, D9891) for three days and changed the media daily. Three days later, the pre-differentiated cells were dissociated with Accutase (STEMCELL Technologies, 07920) and subplated on Poly-L-Ornithine-coated (15 µg/mL; Sigma-Aldrich, P3655) plates with maturation media with doxycycline. Then, the maturation media was changed seven days later by removing half of the media from each well and adding the same amount of fresh media without doxycycline. The differentiated neurons were collected two weeks after differentiation for experiments. The detailed protocol is accessible at the ENCODE portal (https://www.encodeproject.org/documents/d74fb151-366c-4450-9fa0-31cc614035f9/).

### HiC

HiC libraries were generated using the Arima-HiC+ kit (P/N A101020) according to the manufacturer’s protocol. Briefly, differentiated ExNs were fixed in culture plates with 2% PFA (Fisher Scientific F79-500) at room temperature for 10 min. About 0.5 million cells were used as input for each HiC library. The crosslinked chromatin DNA was treated with restriction digestion, biotinylation, proximal ligation. The proximally ligated chromatin was sheared with a Covaris S220 sonicator to fragments between 300 and 1,000 bp. The shared chromatin was then indexed with the Accel-NGS 2S Plus DNA Library Kit (Swift Biosciences, 21024) and amplified with KAPA Library Amplification Kit (Roche, KK2620). The final libraries were purified with AMPure XP beads (Beckman Coulter, A63881) and sequenced on NovaSeq with paired-end 100bp sequencing.

For analyzing HiC data, the 4DN processing pipeline was used, as outlined here: https://data.4dnucleome.org/resources/data-analysis/hi_c-processing-pipeline. In short, reads were mapped to GRCh38 using bwa v0.7.18(45). Valid HiC alignments were filtered using pairtools v1.0.3(46) and returned to a pairs file. Next, biological replicates were merged using https://github.com/4dn-dcic/docker-4dn-hic/blob/master/scripts/run-merge-pairs.sh. Lastly, the pairs files were converted to cool files using cooler v0.9.3(47). Plotted maps from experimental HiC data are at a resolution of 1,000 bp. Each biological replicate was also processed separately and the replicates were visually compared at the variant locus and found to be highly similar.

For contact frequency map visualization, cool files were preprocessed the same as the training datasets for the Akita model(2, 26) to allow for easy visual comparison. Namely, HiC data was normalized with genome-wide iterative correction (ICE)(48). Then, using cooltools v0.7.1, the matrices were smoothed using adaptive coarsegraining, normalized for distance-dependent contact decay. The values were log scaled and limited to (−2,2). Then, again using cooltools v0.7.1, they were linearly interpolated to fill missing values. Lastly, the maps were smoothed using astropy v5.2.2 convolution with a 2D Gaussian filter.

### RNA-Seq

RNA was extracted from two weeks old ExNs differentiated from iPSCs using the RNeasy Mini Kit (Qiagen, 74104). Approximately 4 mg of extracted total RNA was used to prepare libraries for sequencing using the TruSeq Stranded mRNA Library Prep Kit (Illumina, 20020594). Libraries were sequenced on NextSeq 2000 with paired-end 100 bp sequencing.

RNA-sequencing libraries, comprising 6 samples were sequenced to an average depth of 8e7 reads per sample. RNA-Seq reads were aligned to GRCh38.112 using STAR v2.7.11b in gene annotation mode(49). Alignment, RNA-Seq, and Insert Size quality control metrics were generated using Picard v3.1.1. Sample quality was accessed using FASTQC v0.12.1(50) and MultiQC v1.22.2 (51).

Genes with more than 2 counts per million (cpm) in at least 2 of the 6 samples were retained for differential gene expression analysis. Filtered genes were tested for differential expression with DESeq2 v1.44.0(52) and considered significantly differentially expressed with an adjusted p-value < 0.05.

A Gene Ontology (GO)(53) over-representation analysis was performed to test if DEGs from the full deletion are enriched in Biological Process (BP) terms compared to all expressed genes (R package clusterProfiler(54) v4.8.3 and GO.db v3.18.0). Enriched terms were defined by a Benjamini-Hochberg corrected p-value < 0.05 and > 30 DEGs in the category (**SI Appendix, Dataset S2**).

### Data, Materials, and Software Availability

Raw and processed data for HiC and RNA-Seq are available in GEO. SuPreMo V2 can be found on GitHub at https://github.com/ketringjoni/SuPreMo/tree/main.

## Acknowledgements

We thank Ian Jones for guidance with PLAC-Seq data analysis interpretation. We thank Shuzhen Kuang for input on HiC data analysis. We thank Jingjing Li for mentorship and feedback on the project.

## Funding

This work was supported by the NIH 4D Nucleome Project (award #U01HL157989 to K.S.P.), the NIH Office of the Director (award #R03OD034499 to K.S.P.), Additional Ventures, two UCSF Achievement Rewards for College Scientists Scholarships (K.G. and A.E.), Keck Foundation, and Biswas Family Foundation, and Gladstone Institutes.

### List of abbreviations

ISM: in silico mutagenesis
ASD: autism spectrum disorder
TAD: topologically associated domain
CREint: cis-regulatory element interaction
dnSV: de novo structural variant
ExN: excitatory neuron
WT: wild type
DEG: differentially expressed gene

## Author Contributions

K.G. helped conceive the project, performed all computational analyses, other than RNA-Seq, and prepared the manuscript/figures. X.R. performed all wet-lab experiments including CRISPR editing, cell differentiations, and sample preparation for HiC and RNA-Seq. A.E. performed RNA-Seq, differential expression, and GO enrichment analyses. Y.S. helped guide the project and managed the experiments. K.S.P. conceived and managed the project and edited the manuscript. All authors reviewed the manuscript.

## Competing Interest Statement

The authors declare no competing interest.

## Notes

### Competing Interest Statement

The authors have declared no competing interest.

