## Supplemental Information for "De novo structural variants in autism spectrum disorder disrupt distal regulatory interactions of neuronal genes"

**This PDF file includes:**

Figures S1 to S7
Table S1
Legends for Datasets S1 to S2

**Other supporting materials for this manuscript include the following:**

Datasets S1 to S2

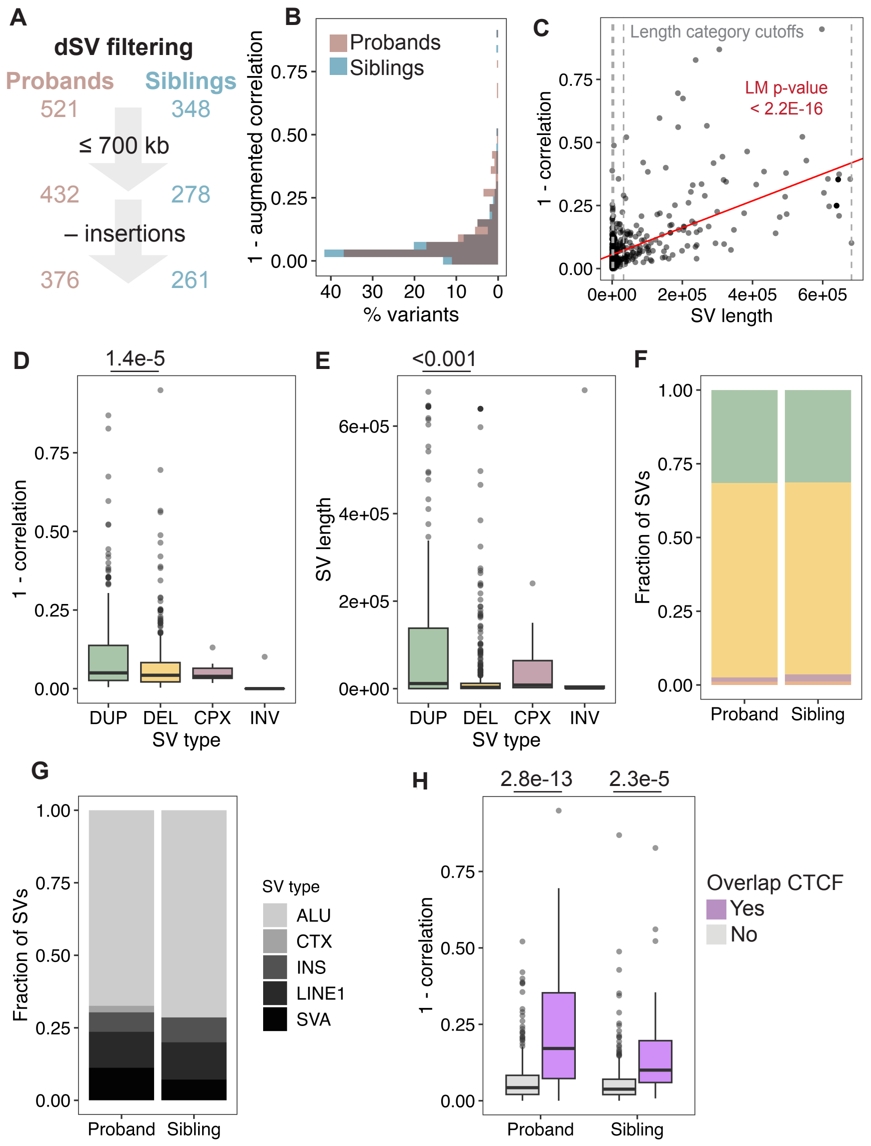


**Fig. S1.** Disruption score trends. A. Graphic of how dnSVs were filtered to ones that are at most ⅔ of the Akita prediction window and types of SVs that are scorable. **B.** Distribution of augmented scores (median score across normal score and three augmentations: 1 bp left shift, 1bp right shift, and reverse complement). Mann-Whitney U test between proband and sibling augmented scores was not significant (p-value = 0.067) but the trend seen in Figure 1B was maintained. **C.** Disruption scores plotted against dnSV length. P-value of linear model predicting scores from length (red line) is shown. Dashed black lines are length cutoffs for the four categories in Figure 1 C and D. **D.** Disruption scores across variant types–duplications (DUP), deletions (DEL), complex variants (CPX), and inversions (INV)–including both proband and sibling dnSVs. **E.** SV length across SV types. **F.** Fraction of dnSV types scored in probands and siblings. For D and E, pair-wise p-value from an ANOVA and TukeyHSD tests were not significant unless the p-value is shown. **G.** Fraction of dnSV types that were excluded from this manuscript because the exact alternate allele sequence is not known: Alu element insertions (ALU), translocations (CTX), insertions (INS), Line1 element insertions (LINE1), and SVA element insertions (SVA). **H.** Comparing disruption scores of dnSVs that do and don’t overlap CTCF binding sites from ExN ChIPseq data in probands and siblings. Mann-Whitney U test p-values shown.

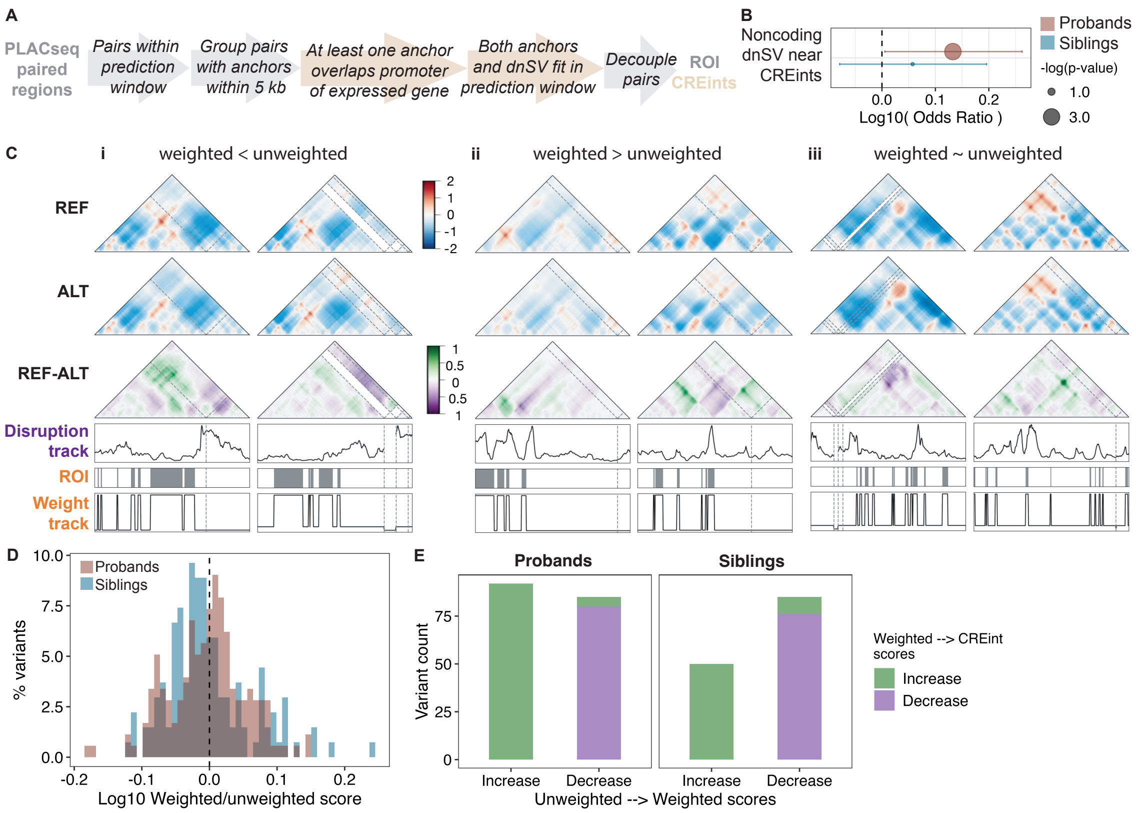


**Fig. S2. Using CREints to generate weighted disruption scores. A.** Pipeline for handling PLACseq paired reads as weights using SuPreMo-Akita (gray) and additional filtering done in the analyses in this paper to generate CREints (beige). **B.** Enrichment of proband and sibling noncoding dnSVs near PLAC-Seq anchors. hg38 was tiled into 1 Mb bins, which were labeled based on whether there encompassed at least one noncoding dnSV (defined as not overlapping Gencode v37 protein coding gene exons) and PLAC-Seq anchor. P-value corresponds to chi-squared test. **C.** Examples of dnSVs that are scored **i.** lower, **ii.** the same, or **iii.** higher when weighing scores by nearby CREints versus not. Akita-predicted contact maps for the reference allele (top maps), alternate allele (middle maps), and difference between the two (bottom maps) are shown alongside disruption track (1 - correlation at each one of 448 bins), region of interest (ROI) track with processed CREint anchors (Methods), and weight track with ROI bins scaled up 10-fold. **D.** Distribution of log fold change between 10-fold weighted scores and unweighted scores in probands and siblings. **E.** Number of variants with weighted scores that are increased or decreased from unweighted scores and CREdit scores that are increased or decreased from weighted scores, separated for probands and sibling dnSVs.

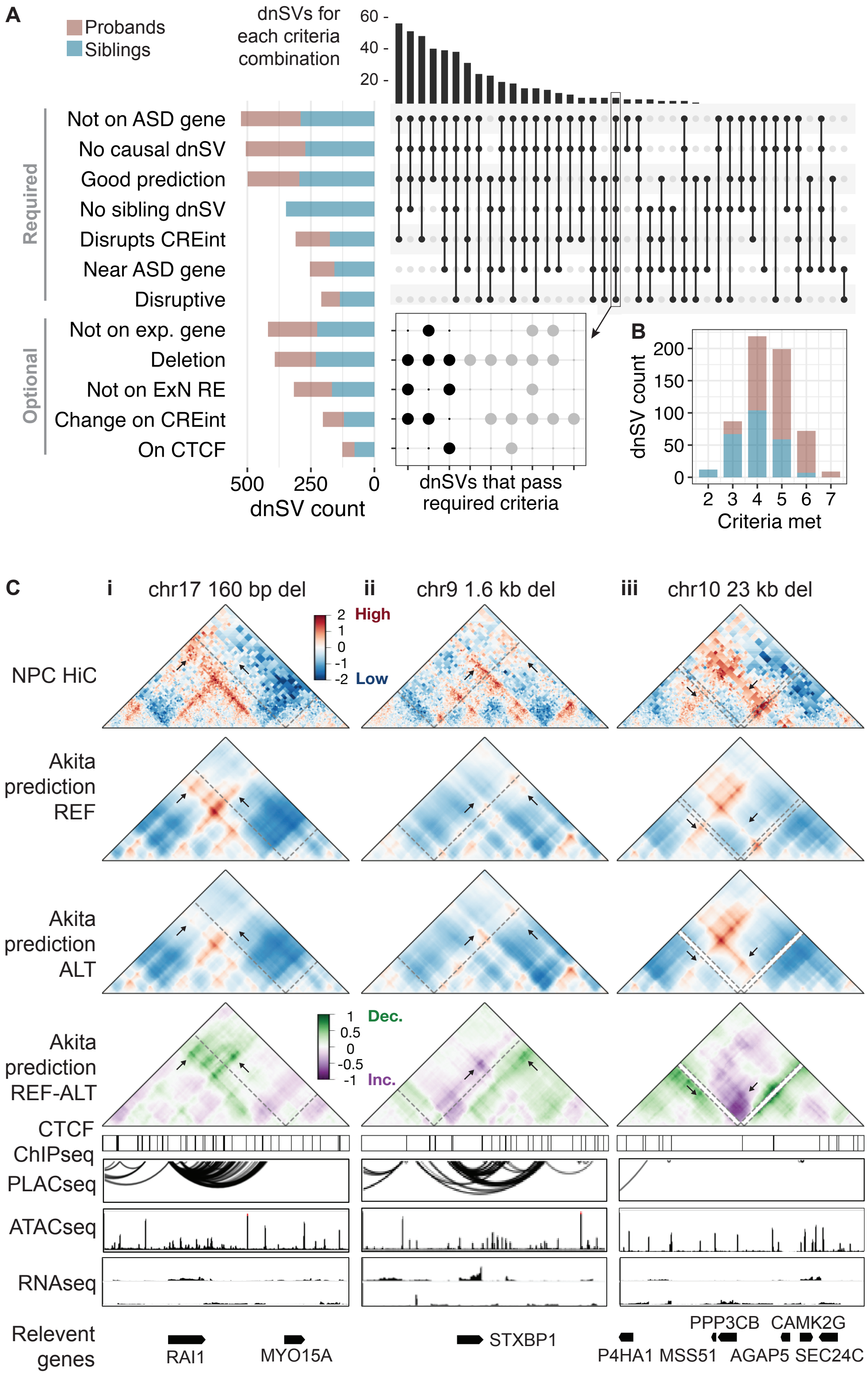
A
Probands
Siblings
dnSVs for each criteria combination
60
40
20
Required
Not on ASD gene
No causal dnSV
Good prediction
No sibling dnSV
Disrupts CREint
Near ASD gene
Disruptive
Optional
Not on exp. gene
Deletion
Not on ExN RE
Change on CREint
On CTCF
500 250 0
dnSV count
dnSVs that pass required criteria
B
dnSV count
200
150
100
50
0
2 3 4 5 6 7
Criteria met
C
i chr17 160 bp del
ii chr9 1.6 kb del
iii chr10 23 kb del
NPC HiC
2 High
1
0
-1
-2 Low
Akita prediction REF
Akita prediction ALT
Akita prediction REF-ALT
1 Dec.
0.5
0
-0.5
-1 Inc.
CTCF ChIPseq
PLACseq
ATACseq
RNAseq
Relevent genes
RAI1
MYO15A
STXBP1
PPP3CB
CAMK2G
P4HA1
MSS51
AGAP5
SEC24C

**Fig. S3. Criteria for prioritizing candidate ASD variants that are likely to be causal via disruption of genomic contacts. A.** Required criteria and distribution of dnSVs that meet them. The 7 criteria are listed on the y axis and the number of proband and sibling dnSVs that meet each is shown in the bar graph. UpSet plot on the right shows the number of dnSVs that meet each combination of the criteria, with 9 dnSVs (boxed) meeting all 7 criteria. These 9 dnSVs are annotated based on the optional criteria they meet in the dot plot below. To define ASD genes, we compiled associated gene sets from three different studies (Methods). **B.** Number of dnSVs that meet different numbers of criteria. **C.** Three candidate dnSVs, i-iii. The contact frequency maps, from top to bottom, correspond to: neural progenitor cell HiC experimental data, Akita prediction for the reference allele, Akita prediction for the alternate allele, and the difference between Akita prediction reference and alternate. Location of the variant in the maps is shown with a dotted gray line. Genes of interest to ASD are shown in black, with arrows showing transcriptional direction. Tracks, from top to bottom, correspond to: iPSC-derived ExNs CTCF ChIPseq and primary ExN H3K4me3 PLACseq loops, ATACseq and RNAseq forward and reverse tracks. Red lines mark the location of the variant.

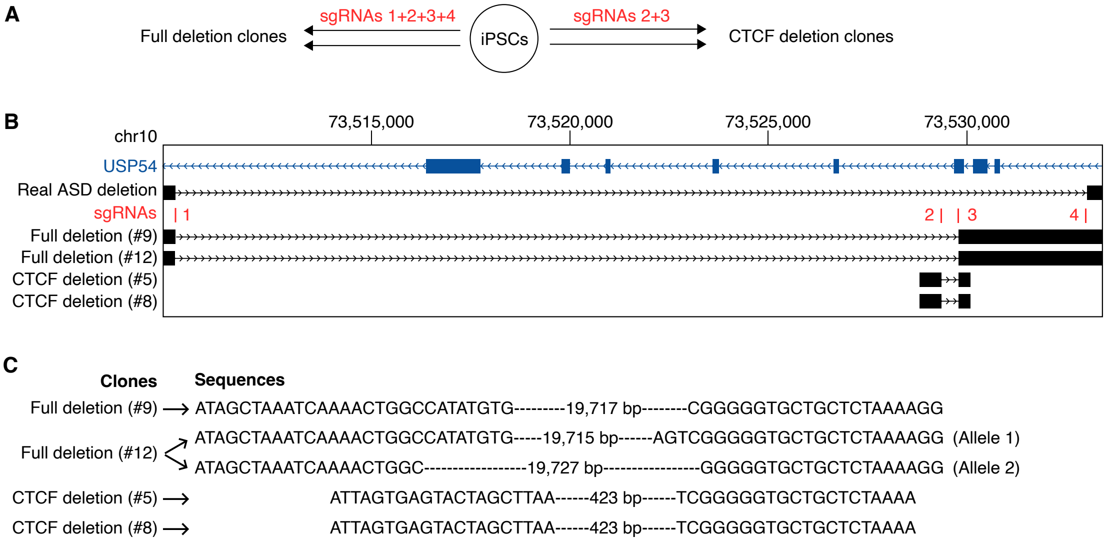


**Fig. S4. Gene editing generated clonal cell lines. A.** Gene editing schematic. iPSCs went through 4 rounds of editing with 2 sets of guides to get two biological replicates of each guide combination. The full deletion clones were generated using guides 1, 2, 3, and 4, and the CTCF deletion clones were generated using guides 2 and 3. **B.** Genomic tracks around the deletion of, from top to bottom, the USP54 gene, the candidate deletion, the 4 gRNAs used, the two full deletion (about 19.7 kb) clones, and the two CTCF deletion (424 bp) clones. **C.** Sequence around the deletion for the 4 clonal cell lines generated. One of the full deletion clonal cell lines (#12) has two different alleles at the deletion site.

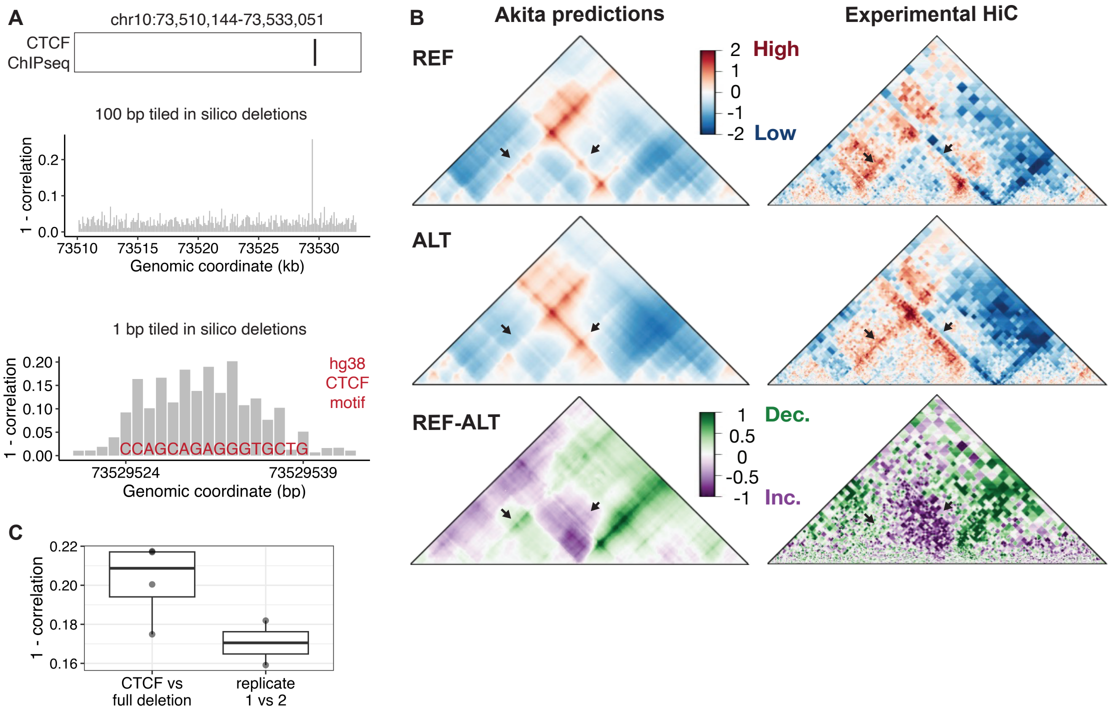


**Fig. S5. Prioritized deletion causes changes to genome folding because of CTCF binding site deletion. A.** Tiled deletions reveal that a CTCF motif is responsible for the predicted change in contact frequency maps. Top: CTCF ChIPseq track in deletion window; middle: disruption scores for 100 bp tiled deletions across the deletion window; bottom: disruption scores for 1 bp tiled deletions across the high scoring 100 bp window reveal a CTCF binding motif in the hg38 reference genome. **B.** Akita-predicted (left) and experimental HiC (right) contact frequency maps for the reference genome (top), CTCF ChIPseq peak deletion (middle) and the difference (bottom). **C.** Boxplot comparing difference between CTCF and full deletion (for all 4 replicate combinations) to difference between within condition replicates.

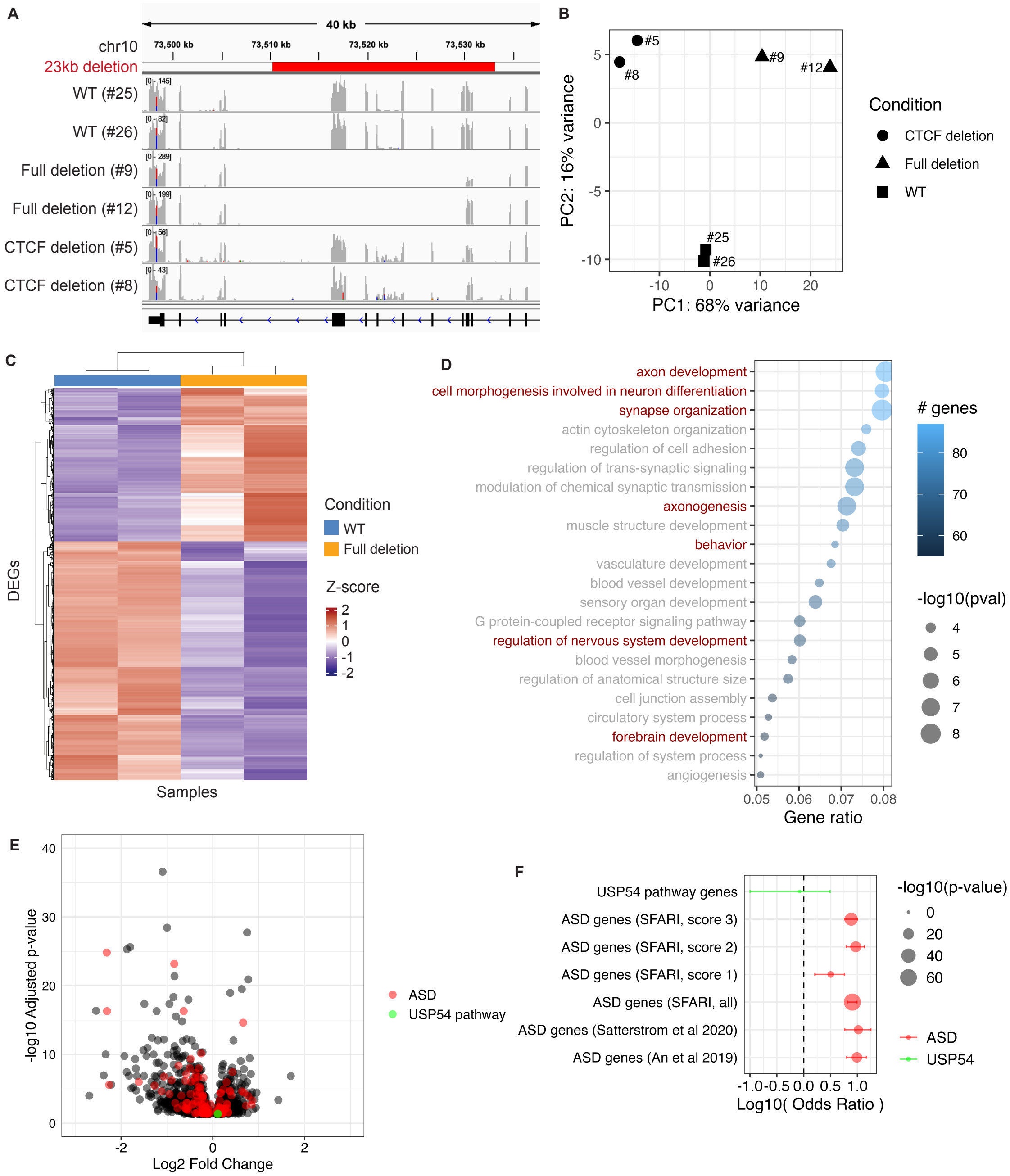
A
40 kb
chr10
73,500 kb
73,510 kb
73,520 kb
73,530 kb
23kb deletion
WT (#25)
WT (#26)
Full deletion (#9)
Full deletion (#12)
CTCF deletion (#5)
CTCF deletion (#8)
B
#5
#8
#9
#12
#25
#26
PC2: 16% variance
PC1: 68% variance
Condition
CTCF deletion
Full deletion
WT
C
DEGs
Samples
Condition
WT
Full deletion
Z-score
D
axon development
cell morphogenesis involved in neuron differentiation
synapse organization
actin cytoskeleton organization
regulation of cell adhesion
regulation of trans-synaptic signaling
modulation of chemical synaptic transmission
axonogenesis
muscle structure development
behavior
vasculature development
blood vessel development
sensory organ development
G protein-coupled receptor signaling pathway
regulation of nervous system development
blood vessel morphogenesis
regulation of anatomical structure size
cell junction assembly
circulatory system process
forebrain development
regulation of system process
angiogenesis
Gene ratio
### genes
-log10(pval)
E
-log10 Adjusted p-value
Log2 Fold Change
ASD
USP54 pathway
F
USP54 pathway genes
ASD genes (SFARI, score 3)
ASD genes (SFARI, score 2)
ASD genes (SFARI, score 1)
ASD genes (SFARI, all)
ASD genes (Satterstrom et al 2020)
ASD genes (An et al 2019)
Log10( Odds Ratio )
-log10(p-value)
ASD
USP54

**Fig. S6. Deletion causes differential expression of ASD and neurodevelopmental genes. A.** IGV Genome browser screenshot of RNAseq bam files for the six cell lines. USP54 transcription continues after the deletion. **B.** PCA plot of the first two principal components shows clustering together of the biological replicates and clear separation between the three different edit conditions. **C.** Heatmap of Z-scores for differentially expressed genes between WT and full deletion clones. Red and purple mark genes that are over- and under- expressed, respectively, in the full deletion cell lines. **D.** GO enrichment terms for DE genes in full deletion cell lines. Terms related to development or, specifically, neurodevelopment are highlighted in orange and red, respectively. Background is genes with more than 2 counts per million in at least 2 of the 6 samples. GO terms with >= 50 genes, adjusted p-value < 0.001, and gene ratio > 0.05 are shown here. All GO terms are included in Supplementary Table 3. **E.** Volcano plot showing DEGs in full deletion clones, with labels showing ASD genes and genes that are in the Ubiquitin-Proteasome Dependent Proteolysis pathway, which USP54 is in. Axes are cut off for easier visualization and exclude two genes. **F.** Enrichment of gene sets (ASD and USP54 pathway genes) in DEGs.

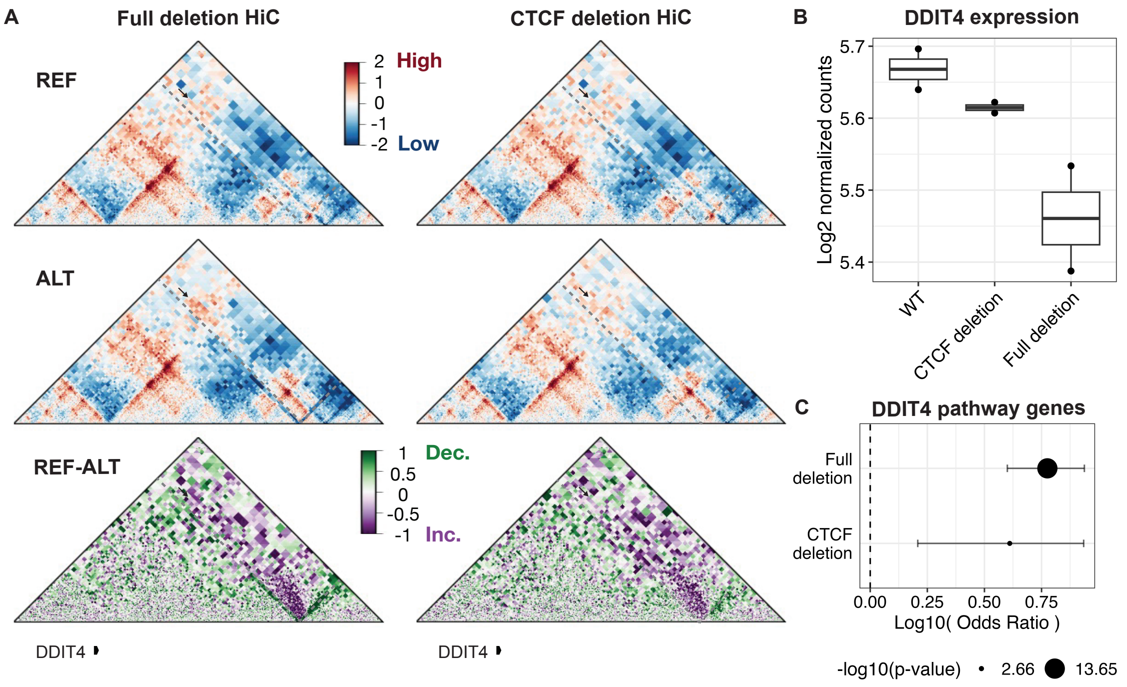


**Fig. S7. Changed genomic contacts could partially contribute to differential expression of DDIT4. A.** Experimental HiC contact frequency maps for the full deletion (left) and CTCF deletion (right) in a X kb region around the deletion (dotted gray line) and DDIT4 (noted below maps). Changes caused by the deletions on DDIT4 contacts are marked with arrows. **B.** Normalized RNAseq counts for DDIT4 in all three editing conditions for both biological replicates. **C.** Enrichment of genes from the PI3K/AKT/mTOR (PAM) pathway, which DDIT4 is a part of, in DEGs.

**Table S1. Oligo sequences for creating gene edited cell lines.**

| Type | Oligo | Sequence |
|---|---|---|
| sgRNA | sgRNA1 | AAAACTGGCCATATGTGTGG |
| sgRNA | sgRNA2 | GATGATTCCTTAAAACTGAT |
| sgRNA | sgRNA3 | CCATTCGCTGGATTAGTCGG |
| sgRNA | sgRNA4 | GTGAGTACTAGCTTAATATG |
| Genotyping primers | Full_deletion_F | CTGAGAGATACATGCCAAATTATTTATG |
| Genotyping primers | Full_deletion_R | TGGTGGTTTATAAGCTGTGGGG |
| Genotyping primers | CTCF_deletion_F | CGTGAGCCAACATGCCTGACC |
| Genotyping primers | CTCF_deletion_R | CATCTGGCAAGGGGCACTTTTACTC |
| Genotyping primers | Full_deletion_allele_C_F | CAACATGGCAAAACCCTGTC |
| Genotyping primers | Full_deletion_allele_T_F | CAACATGGCAAAACCCTGTT |
| In vitro transcription gRNAs | sgRNA_1F | TAATACGACTCACTATAGAAAACTGGCCATATGTGTGG |
| In vitro transcription gRNAs | sgRNA_2F | TAATACGACTCACTATAGGATGATTCCTTAAAACTGAT |
| In vitro transcription gRNAs | sgRNA_3F | TAATACGACTCACTATAGCCATTCGCTGGATTAGTCGG |
| In vitro transcription gRNAs | sgRNA_4F | TAATACGACTCACTATAGGTGAGTACTAGCTTAATATG |
| In vitro transcription gRNAs | sgRNA_1R | TTCTAGCTCTAAAACCCACACATATGGCCAGTTTT |
| In vitro transcription gRNAs | sgRNA_2R | TTCTAGCTCTAAAACATCAGTTTTAAGGAATCATC |
| In vitro transcription gRNAs | sgRNA_3R | TTCTAGCTCTAAAACCCGACTAATCCAGCGAATGG |
| In vitro transcription gRNAs | sgRNA_4R | TTCTAGCTCTAAAACCATATTAAGCTAGTACTCAC |

**Dataset S1 (separate file).** Filtered genes (more than 2 cpm in at least 2 samples) and their normalized expression levels across all six cell lines. The last four columns provide the adjusted p values and log fold change of DEGs.

**Dataset S2 (separate file).** GO biological processes for the 1,102 DEGs in the full deletion cell line, and accompanying statistics.
